# Comparison of the 3-D patterns of the parasympathetic nervous system in the lung at late developmental stages between mouse and chicken

**DOI:** 10.1101/318113

**Authors:** Tadayoshi Watanabe, Ryo Nakamura, Yuta Takase, Etsuo A. Susaki, Hiroki R. Ueda, Ryosuke Tadokoro, Yoshiko Takahashi

## Abstract

Although the basic schema of the body plan is similar among different species of amniotes (mammals, birds, and reptiles), the lung is an exception. Here, anatomy and physiology are considerably different, particularly between mammals and birds. In mammals, inhaled and exhaled airs mix in the airways, whereas in birds the inspired air flows unidirectionally without mixing with the expired air. This bird-specific respiration system is enabled by the complex tubular structures called parabronchi where gas exchange takes place, and also by the bellow-like air sacs appended to the main part of the lung. That the lung is predominantly governed by the parasympathetic nervous system has been shown mostly by physiological studies in mammals. However, how the parasympathetic nervous system in the lung is established during late development has largely been unexplored both in mammals and birds. In this study, by combining immunocytochemistry, the tissue-clearing CUBIC method, and ink-injection to airways, we have visualized the 3-D distribution patterns of parasympathetic nerves and ganglia in the lung at late developmental stages of mice and chickens. These patterns were further compared between these species, and three prominent similarities emerged: (1) parasympathetic postganglionic fibers and ganglia are widely distributed in the lung covering the proximal and distal portions, (2) the gas exchange units, alveoli in mice and parabronchi in chickens, are devoid of parasympathetic nerves, (3) parasympathetic nerves are in close association with smooth muscle cells, particularly at the base of the gas exchange units. These observations suggest that despite gross differences in anatomy, the basic mechanisms underlying parasympathetic control of smooth muscles and gas exchange might be conserved between mammals and birds.

**Highlights:**

- 3-D patterns of parasympathetic nerves are visualized in mouse and chicken lungs.

Comparison of these patterns reveals three prominent similarities between mouse and chicken:

1. VAChT-positive postganglionic fibers and ganglia are widely distributed in the lung.
2. Gas exchange units are devoid of parasympathetic nerves.
3. Parasympathetic nerves are in close association with smooth muscle cells.

## Introduction

It is increasingly appreciated that the molecular and cellular mechanisms underlying early development are largely shared among different species in vertebrates. This also holds true for the mechanisms of organogenesis and their physiology, which are particularly highly shared among amniotes (mammals, birds, reptiles). However, there are some exceptions, and the lung is one such example. Although the extensively branched airways in the lung offer a platform for gas exchange between O_2_ and CO_2_ in mammals and birds, the anatomical structures of the lung are considerably different between them (Schachner et al., 2013; Lambertz et al., 2015; Cieri and Farmer, 2016). Unlike in mammals, where inhaled and exhaled gases mix in the lungs, in birds, the air flows unidirectionally at inspiration and expiration. This provides birds with higher efficiencies of gas exchange than mammals (Barnas et al., 1978; Banzett et al., 1991; Brown et al., 1997; Boggs et al., 1998; Nasu, 2005; Çevik-demirkan et al., 2006; Reese et al., 2006; West et al., 2007; Maina, 2008; Plummer and Goller, 2008; Makanya and Djonov, 2009; Maina, 2015, 2017). This high efficiency of gas exchange, together with the avian-specific bellow-like air sacs appended to the lung, enables birds to fly at high altitudes and/or long distances. Such remarkable differences between avian and mammalian lungs have long attracted investigators in many biological fields including developmental biology and locomotive biology. However, besides anatomy and some physiology (Boggs et al., 1998), comparative studies of the lung differences at the cellular neurological level remains poorly explored.

Like other organs in the body, the lung physiology is governed by the autonomic nervous system, which is composed of sympathetic- and parasympathetic neurons (McCorry, 2007; Wehrwein et al., 2016; Karemaker, 2017). Previous studies, predominantly in mammals, reveal that the parasympathetic nervous system is particularly important in lung physiology, whereas the sympathetic nervous system plays less important roles (Mazzone and Canning, 2013). It has been proposed that the parasympathetic control of the airway constriction (bronchoconstriction) is mediated by acetylcholine and smooth muscles (Sparrow et al., 1994; Sparrow et al., 1995). In pathological conditions such as asthma, a common disease in the lung, a denervation of parasympathetic-containing nerves ameliorates hyper-reactivity caused by increased constrictive tone (Balogh et al., 1957; Lewis et al., 2006; Liu et al., 2014). Thus, parasympathetic nervous system plays important roles in the contraction of bronchial tubes both in normal and pathological conditions.

The initiation of this parasympathetic innervation during development has been studied. In mice, immunostaining analyses in histological sections and whole-mounted specimens showed that parasympathetic neurons start to innervate the lung by embryonic day 12.5 (E12.5), where they are in close association of growing bronchial structures (Lath et al., 2012). In chickens, precursors of parasympathetic ganglia originating from vagal region-derived neural crest cells reach the lung buds by E5 (Burns and Delalande, 2005). However, it remained unknown how the global innervation patterns of the parasympathetic neurons in the lung are established at late stages during development both in chickens and mice, since the branches and their patterning become progressively thick and complex, hampering 3-D visualization of airways and nerves by conventional methods.

In this study, we investigated the distribution patterns of the parasympathetic nervous system in whole-mounted lung specimens with a particular focus on late embryonic and newborn stages, and we compared the patterns between mice and chickens. Two technical breakthroughs were undertaken: one was to employ the advanced tissue clearing method (Susaki et al., 2014; Tainaka et al., 2014; Susaki et al., 2015), which allows high-resolution 3-D visualization of the intricate innervation patterns in the lung. The other was immunocytochemistry using recently raised antibody against chicken vesicular acetylcholine transporter (VAChT) protein, which distinguishes parasympathetic neurons from sympathetic postganglionic neurons (Watanabe et al., 2017).

In both species, parasympathetic nerves are in close association with smooth muscles along the bronchial tubes, and they innervate widely in the lung to the periphery. Importantly, the gas exchange tissues, alveoli in mice and atria (air capillaries) in chickens, are devoid of the parasympathetic innervation. Thus, despite considerable differences in anatomy, mice and chickens appear to employ similar mechanisms for the neurophysiology of gas exchange. Some of the data photographs shown in the Figures are images extracted from 3-D movies, 9 of which are appended as Supplementary Materials.

## Materials and Methods

### Experimental animals

Jcl:ICR (ICR) strain (CLEA Japan, Inc., Tokyo, Japan) mice and fertilized eggs of the Hypeco nera chicken strain were purchased from Shimizu Laboratory Supplies Co., Ltd. (Kyoto, Japan) and Shiroyama poultry farm (Kanagawa, Japan), respectively.

### Visualization of lung airways by highlighter ink

Following the dissection of the lung, commercially available highlighter ink (Takase et al., 2013) was injected into the trachea, followed by fixation in 4% (w/v) paraformaldehyde (PFA)/phosphate buffered saline (PBS: 0.14 M NaCl, 2.7 mM KCl, 10 mM Na_2_HPO_4_-12H_2_O, 1.8 mM KH_2_PO_4_).

### Tissue clearing by CUBIC

Tissue clearing was basically performed according to (Susaki et al., 2014; Tainaka et al., 2014; Susaki et al., 2015) with following modifications: fixed specimens were immersed in CUBIC-reagent 1 and CUBIC-reagent 2 consecutively, followed by whole-mount immunostaining. The specimens were immersed again in CUBIC-reagent 2 because this process declined an excess amount of staining (noise) with specific signals retained. When combined with ink injection, this was implemented after the immunostaining, and the following CUBIC-reagent 2 treatment was shorter than 30 minutes.

### Immunohistochemistry with the mouse lung

Two kinds of anti-VAChT antibodies were used to stain mouse tissues: one was purchased from Synaptic System (139105, 1:500), and the other from Abcam (ab62140, 1:200), for which two different fixation protocols were employed: the fixation in PFA and in the solution of dimethylsulfoxide (DMSO): methanol, respectively.

#### Staining with PFA-fixed specimens

Lungs were fixed in 4 % PFA/PBS at 4 °C with gentle shaking overnight, and washed in PBS three times for 1 hour each. For the staining for VAChT and Tuj1, 300-400 μm sections of lung lobes were obtained using a micro-slicer (SUPER MICRO SLICER ZERO 1, DOSAKA EM). Following the CUBIC-tissue clearing, the slices were treated with 0.3 % H_2_O_2_ in PBS for 1 hour and washed in 0.1 % Triton X-100 in PBS three times for 1 hour. After pre-blocking with blocking solution (0.5 % BR/PBST; 0.5 % Blocking reagent [BR; Roche, 11096176001]/PBST [0.5 % TritonX-100]) twice for 1 hour each, the slices were incubated for 48 hrs at 4 °C with primary antibodies, anti-VAChT (1:500, Synaptic System, 139105) and anti-Tuj1 (1:300, R&D systems, MAB1195), in 0.5 % BR/PBST. The slices were washed 5 times for 1 hour each in 0.5 % BR/PBST at room temperature, and incubated with secondary antibodies conjugated with anti-mouse IgG-Alexa 488 and anti-rabbit IgG-Alexa 568 (1:500, Molecular probes) for 48 hrs periods at 4 °C. After washing 5 times in 0.1 % (v/v) Tween 20 in PBS, the slices were reacted with 1 μg/ml DAPI (Nacalai Tesque, 11034-56) in PBST (0.1 % Tween 20), and washed in PBST (0.1 % Tween 20) three times for 30 minutes each.

#### Staining with DMSO:methanol-fixed specimens

Lungs were fixed in DMSO:methanol (1:4) at 4 °C with gentle shaking for overnight. Fixed lungs were then incubated in 30 % H_2_O_2_:DMSO:methanol (1:1:4) for 8 hours at room temperature with gentle rocking, and stored at -20 °C in 100 % methanol. Whole mount immunohistochemistry was performed as previously described (Metzger et al., 2008) with the following modifications: specimens were incubated with anti-VAChT (1:200, Abcam, ab62140) diluted in 0.5 % BR/PBST for 2 or 3 nights at 4 °C, and subsequently with secondary antibodies conjugated with anti-rabbit IgG-Alexa 488, anti-mouse IgG-Alexa 647 (1:500, Molecular Probes), and AmpliStain^TM^ anti-Goat 1-Step HRP (1:500, Stereospecific Detection Technologies, AS-G1-HRP) overnight at 4 °C. The specimens were reacted with 1:200 dilution of Cy3-tyramid (Perkin–Elmer, FP1170) in 1× Amplification diluent (Perkin–Elmer, FP1135) for 1 hour at room temperature, and washed three times in TNT (0.1 M Tris-HCl [pH7.5], 150 mM NaCl, 0.1 % Tween 20) to terminate the reaction. Anti-PGP9.5 (1:200, Invitrogen, 38-1000) and anti-Tuj1 (1:300, R&D systems, MAB1195) were also used for tissues fixed in DMSO:methanol.

### Immunohistochemistry with chicken embryos

Lungs fixed in 4 % PFA/PBS overnight were washed in 0.1 % Triton X-100 in PBS (PBST) three times for 5 minutes each, and processed to the CUBIC-tissue clearing. Subsequently, the specimens were treated with 0.3 % H_2_O_2_ in methanol for 1 hour and washed in PBST three times for 5 minutes each. After 1 hour of pre-blocking with 1 % BR in PBST, the specimens were incubated overnight at 4 °C with primary antibodies in 1 % BR/PBST, washed three times in PBST and incubated with 1:1000 dilution of anti-mouse or anti-rabbit peroxidase polymer (Dako), and with 1:400 dilution of anti-mouse IgG-Alexa 488-conjugated antibody (Molecular Probes) with DAPI for 30 minutes. After washing six times in TNT, the lungs were reacted with 1:200 dilution of Cy3-tyramid in 1× Amplification diluent for 5 minutes at room temperature. The reaction was terminated by washing three times in TNT. The primary antibodies were mouse monoclonal- and rabbit polyclonal antibodies against chicken VAChT (1:200, Watanabe et al., 2017), mouse anti-acetylated tubulin (1:200, SIGMA-ALDRICH, T6793), and mouse anti-αSMA (1:300, SIGMA-ALDRICH, A5228).

### Immunohistochemistry with frozen sections

Frozen sections (80 μm for mice, 40 μm for chickens) of PFA-fixed tissues were prepared using a cryostat (MICROM, HM500 OM). The sections were washed in PBST (0.1 % Triton X-100) three times for 30 minutes, treated with 0.3 % H_2_O_2_ in PBS for 30 minutes, rinsed in PBST three times for 30 minutes, and permeabilized with 1.5 % TritonX-100 for 10 minutes. The sections were washed in PBTD (0.15 % TritonX-100, 5 % DMSO in PBS), and blocked with 0.5 % BR in PBTD for 1 hour at room temperature. They were subsequently incubated with primary antibodies, anti-VAChT (1:400, Synaptic System, 139105), anti-Tuj1 (1:300, R&D systems, MAB1195), anti-αSMA (1:300, SIGMA-ALDRICH, A5228), and anti-Tyrosine hydroxylase (TH) (1:200 or 1:400, Pel-Freez, P40101-150; 1:200, Millipore, MAB318) for overnight at 4 °C, washed in PBST three times for 30 minutes, and pre-blocked with the 0.5 % BR in PBTD for 1 hour. They were incubated with the secondary antibodies conjugated with anti-mouse or rabbit IgG-Alexa 488 and anti-rabbit or mouse IgG-Alexa 568 (1:500, donkey; Molecular Probes) for overnight at 4 °C. For TH staining with chicken embryos, the sections were reacted with 1:200 dilution of Cy3-tyramid in 1× Amplification diluent for 5 minutes at room temperature after washing six times in TNT. The reaction was terminated by washing three times in TNT. They were washed in 0.1 % Tween 20 in PBS three times for 30 minutes, stained with 1 μg/ml DAPI in 0.1 % Tween 20 in PBS for 10 minutes at room temperature, and mounted with Fluoromount (Diagnostic BioSystems, K 024).

### Histology

Hematoxilyn-Eosin (HE) staining: frozen sections of 10 μm were stained in Mayer’s Hematoxylin (Muto Pure Chemicals CO., LTD, 3000-2) for 10 seconds, and washed in running tap water. Following a replacement of ethanol solution, the sections were stained in Eosin Alcohol Solution (Wako, 050-06041) for 5 minutes. After dehydration in an ethanol series, the sections were cleared in xylene and mounted.

### Microscopy

Images of ink-injected lungs were obtained using Leica MZ 10F. The immuno-stained tissues and sections were imaged with the Nikon A1R confocal microscope, or by Zeiss Imager Z1 microscope with ApoTome.

## Results

### Overview of the lung anatomy in mice, and visualization of airways by ink injection

In the lung of mice, the trachea undergoes progressive branching that produces numerous terminal bronchioles (Fig. 1A). The tip of each bronchiole is a dead-end with alveoli, where gas exchange takes place. The lung volume substantially changes during respiration.

**Figure 1.**
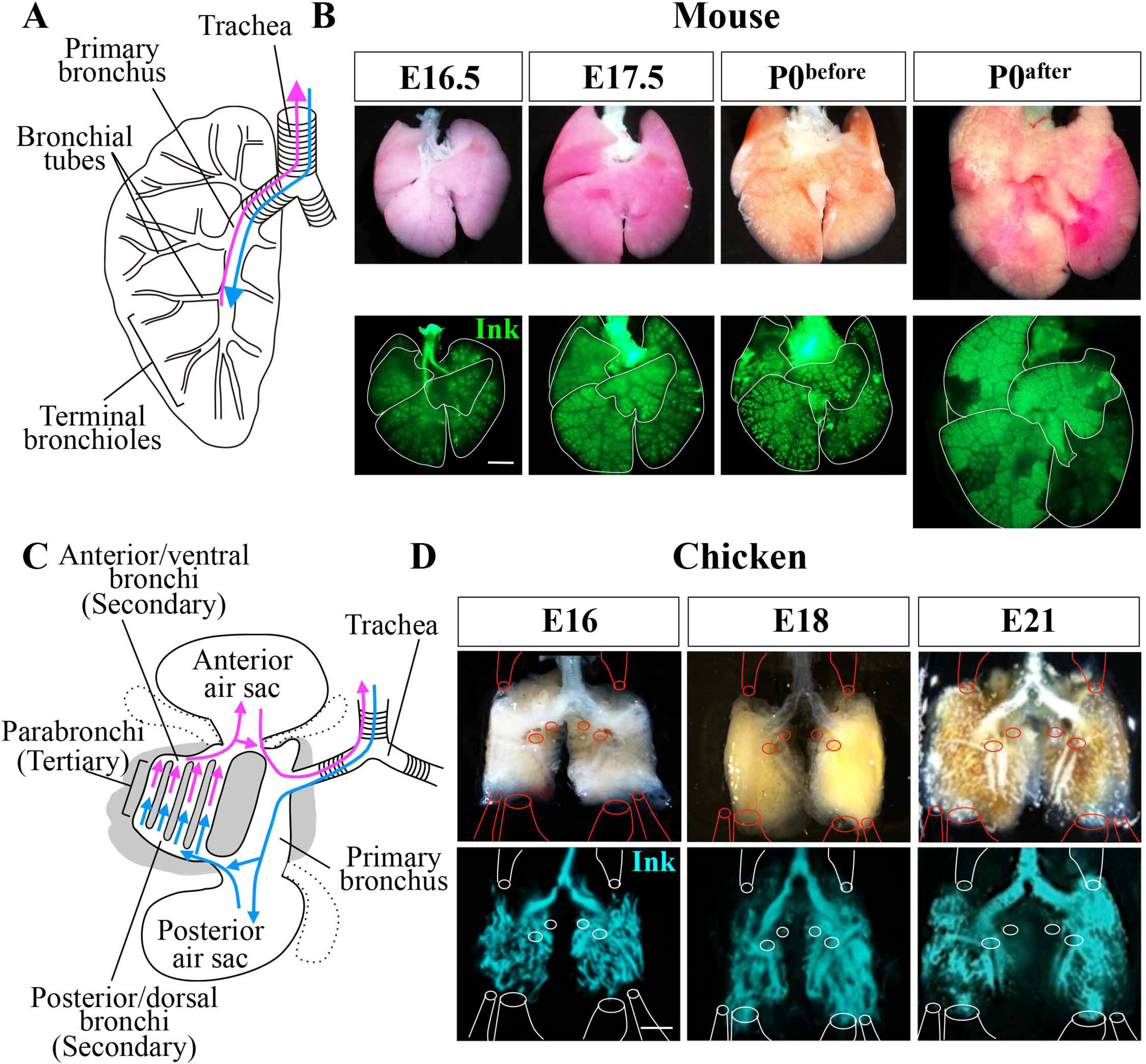
Gross anatomy and visualization of airways by ink injection in the developing lung of mice and chickens. (A, B) Mouse lungs. (C, D) Chicken lungs. (A, C) Schematic diagram showing gross anatomy and airflows. (B, D) Branching patterns in developing lungs visualized by highlighter ink injection at the stages indicated. Solid lines enclose different lobes in (B), and indicate the positions of connecting points of the air sacs (D). Upper panels are bright fields, and lower panels are images captured by a GFP-filter. Scale bars: 1 mm in (B), 2 mm in (D). n=3 per each experiment.

In this study, we examined the distribution patterns of parasympathetic neurons in the developing lung in relation to the morphogenetic patterns of airway branching at relatively late developmental stages from E16.5 through the birth. Since the airway branches are very complex at these stages, the overall structures of the airways, including their peripheral bronchioles, were visualized by injecting fluorescent ink, reported to be useful to visualize embryo-wide vascular network (Takase et al., 2013). The pulmonary branching pattern started to become elaborate by E16.5, and this elaboration progressed through the birth. Embryos just before birth (P0^before^) and after birth (P0^after^) were siblings of the same mother, which were harvested when the first and second pups were delivered. Increased volume of the lung at P0^after^ compared to P0^before^ was evident due to the first breath after delivery (Fig. 1B).

### Overview of the lung anatomy in chickens, and visualization of airways by ink injection and tissue clearing

In the lung of chickens, the trachea branches into a pair of primary bronchi, which connects the secondary bronchi, posterior/dorsal (hereafter called “posterior”) and anterior/ventral (hereafter called “anterior”) bronchi (Brown et al., 1997; Cieri and Farmer, 2016). Between the posterior and anterior bronchi, numerous air tubes, called parabronchi, are arranged in parallel with each other, where gas exchange takes place. Five air sacs composed of 3 anterior and 2 posterior are appended to each of the right and left lobes of the lung (Fig. 1C; hereafter the diagram of the chicken lung displays only two air sacs for simplicity) (Kitazawa et al., 1976; Sakiyama et al., 2000; Maina, 2003). After the primary bronchus, the inspired air flows into the posterior air sacs, from which the air goes anteriorly through parabronchi into anterior bronchi and/or anterior air sacs. The CO_2_-rich air finally returns to the trachea and is expired from the body. The main part of the lung filled with parabronchi does not change in volume during respiration, whereas air sacs serve as bellows that expand and compress dynamically.

In ink-injected and tissue-cleared lungs, the primary bronchi, the posterior and anterior bronchi, and the parabronchi were visualized. During the preparation of specimens, the air sacs, which were extremely thin and fragile, were removed from the main part of the lung. In Fig. 1D, the connecting positions of air sacs are depicted. Although the very complex structures of parabronchi were not easily recognized in low magnification (Fig. 1D), the fine structures of air capillaries/atria within each parabronchus were successfully visualized by this method in higher magnification as explained in details below (for example Fig. 5). The complexity in the parabronchial structures was already seen at E16, and its overall structure was basically maintained until hatching (E21).

### Global distribution patterns of VAChT-positive nerves and ganglia in the lung of mouse late embryos

It has previously been reported that parasympathetic nerves start to innervate the developing bronchi in the lung of mouse embryos by E12.5 (Lath et al., 2012), when elaborate arborization is not yet obvious. To understand the global distribution of parasympathetic nerves in relation to the late bronchial patterns, the lung of E16-17 mouse embryos was stained in whole-mount with the anti-VAChT antibody (commercially available, see Materials and Methods), followed by tissue clearing by the CUBIC method. Since the VAChT-staining yielded some background signals, specimens were co-stained with the pan-neural marker Tuj1, which recognizes neuronal class III β-Tubulin.

Around the primary bronchus just after the first branching of the trachea at E16, VAChT-positive ganglia, recognized as a cohort of neuronal cells, were densely located. The size of these ganglia was relatively large up to 194 μm in diameter (Fig. 2A-C, F, G). Although VAChT-staining does not distinguish between presynaptic (vagal) neurons and post-synaptic (neural crest-derived) neurons, these large ganglia are presumably the sites where the pre- and post-ganglionic neurons synapse. VAChT-positive nerve fibers were also densely located around the thick airways, and these fibers extended peripherally in a progressively scarce manner along bronchial tubes seen at E16 (Fig. 2A and D). Some neurons/ganglia were VAChT- and Tuj1-double positive, whereas Tuj1-single positive neurons, presumably sensory or sympathetic neurons, were also observed (Fig. 2D, see also below for sympathetic neurons).

**Figure 2.**
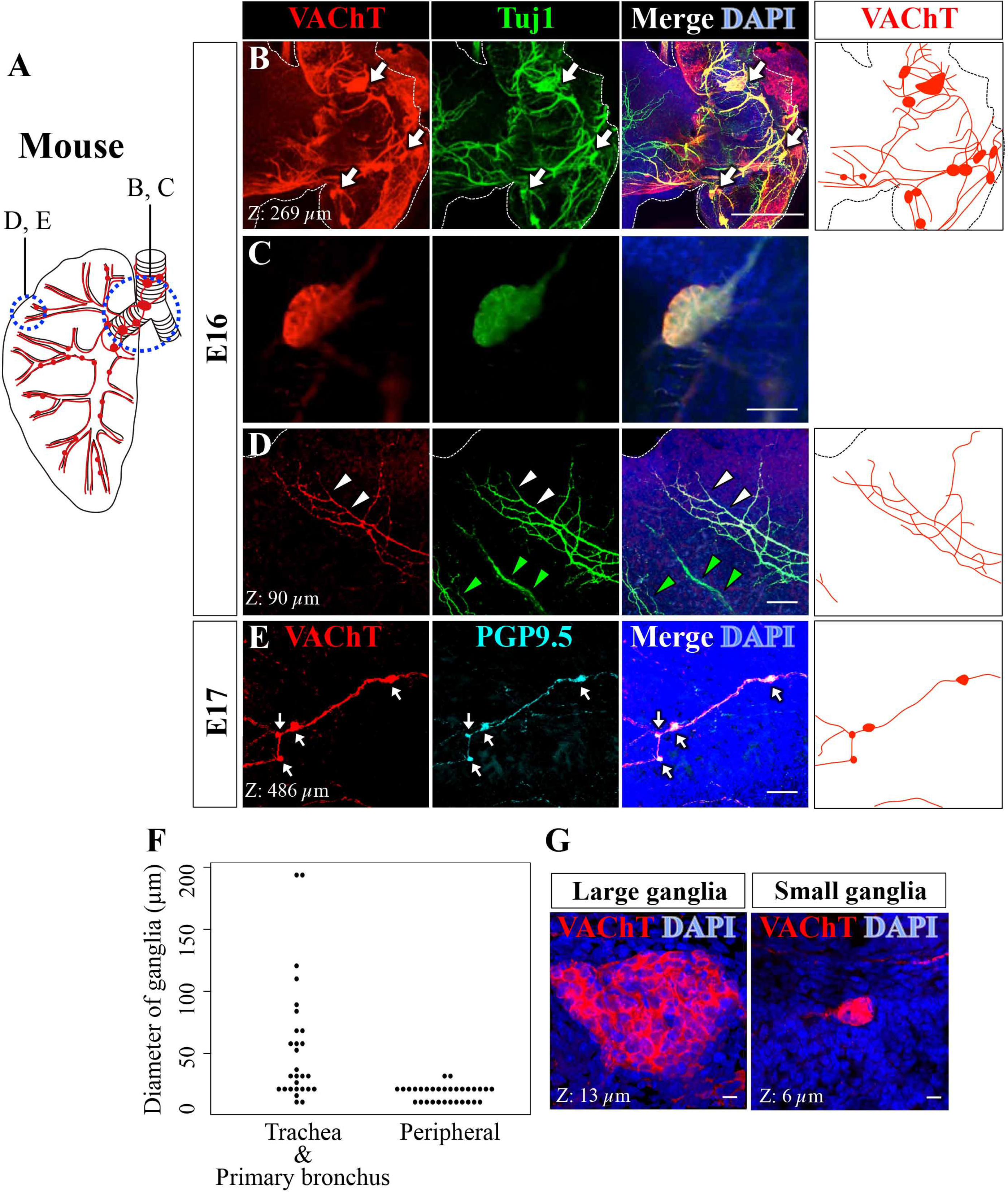
Global distribution patterns of VAChT-positive nerves and ganglia in the mouse lung. (A) Diagram displaying the positions of the images (B-E), and a summary of the distribution patterns of VAChT-positive nerves and ganglia. (B-D) Confocal projection images of a mouse lung lobe at E16, staining for VAChT (red), Tuj1 (green), and DAPI (blue). (B, C) VAChT signals in the primary bronchial tube near the trachea (B), and a ganglion was magnified in (C). (D) VAChT-positive nerves at the periphery. (E) Nerves at the periphery of the lung at E17 stained for VAChT (red), PGP9.5 (cyan), and DAPI (blue). n=3 per each experiment. (F) Site-specific size of VAChT-positive ganglia in the mouse lung at E18. n=4. (G) Confocal projection images of large and small ganglia stained for VAChT (red) and DAPI (blue). White arrows and arrowheads indicate VAChT-positive ganglia and nerves, respectively. Green arrowhead indicates Tuj1-single positive nerves. Dashed lines indicate margins of lung and trachea. Scale bars: 500 μm in (B), 50 μm in (C), 100 μm in (D, E). 10 μm in (G). Confocal images were obtained at 3.20 μm intervals in (B-E), 0.57 μm intervals in (G), for Z-projections as indicated.

At the periphery, large ganglia were not found. Instead, small ganglia of 10 μm to 30 μm in diameter were localized along VAChT-positive nerves (Fig. 2E-G). These small ganglia were explicitly visualized by staining with anti-protein gene product 9.5 (PGP9.5) antibody, which preferentially stains cell bodies of neurons (Fig. 2E) (Sparrow et al., 1999; Tollet et al., 2001). Both the sparse and small ganglia along the peripherally extending nerves and the dense and large ganglia around the primary bronchus were observed in postnatal day 1 (P1) pups, the latest stage examined in this study (Suppl. Fig. 1A, B). This is the first demonstration of the small parasympathetic ganglia at the periphery of the lung at late stages of mouse development.

### Global distribution patterns of VAChT-positive nerves and ganglia in the lung of chicken late embryos

Because of the highly complicated anatomy of the chicken lung at late stages, we combined three methods to clarify the relative positions of VAChT-positive nerves and airways to visualize the distribution pattern of parasympathetic nerves in the whole-mount chicken lung: one was immunocytochemistry using the recently raised antibody against the chicken VAChT protein (Watanabe et al., 2017), second was CUBIC tissue-clearing method, and the third was delineation of airways by injecting highlighter ink, although the ink rapidly diffused in a CUBIC solution (see Materials and Methods). These combined techniques allowed us to locate with high resolution the parasympathetic nerves and ganglia in regard to the positions of airways.

As shown in Fig. 3A-D and also in movies 1-3, in the lung of E16 chicken embryos, VAChT-positive ganglia were located at branching points from secondary (anterior and posterior) bronchi to parabronchi. We also found VAChT-positive ganglia at transition points between primary and secondary bronchi (Fig. 3A, E). Essentially, a single ganglion was present at each branching point (Fig. 3, A-D). Similarly to the mouse, relatively large (57 μm to 171 μm in diameter) and small (5 μm to 105 μm) ganglia were seen in proximal and distal regions in the lung, respectively (Fig. 3D, E, G). VAChT-positive nerves were observed both outside and inside parabronchi. The nerves within the parabronchi were a mesh-like structure with no accompanying ganglia (Fig. 3A-C), which is shown in more details below (see also Fig 5). More proximally toward the branching point of primary bronchus to the trachea, thicker bundles of VAChT-positive nerves were observed, which we presume to be presynaptic vagal nerves (Fig. 3A, F). In the air sacs, VAChT-positive nerves were sparsely present with no recognizable ganglia (Suppl. Fig. 2).

**Figure 3.**
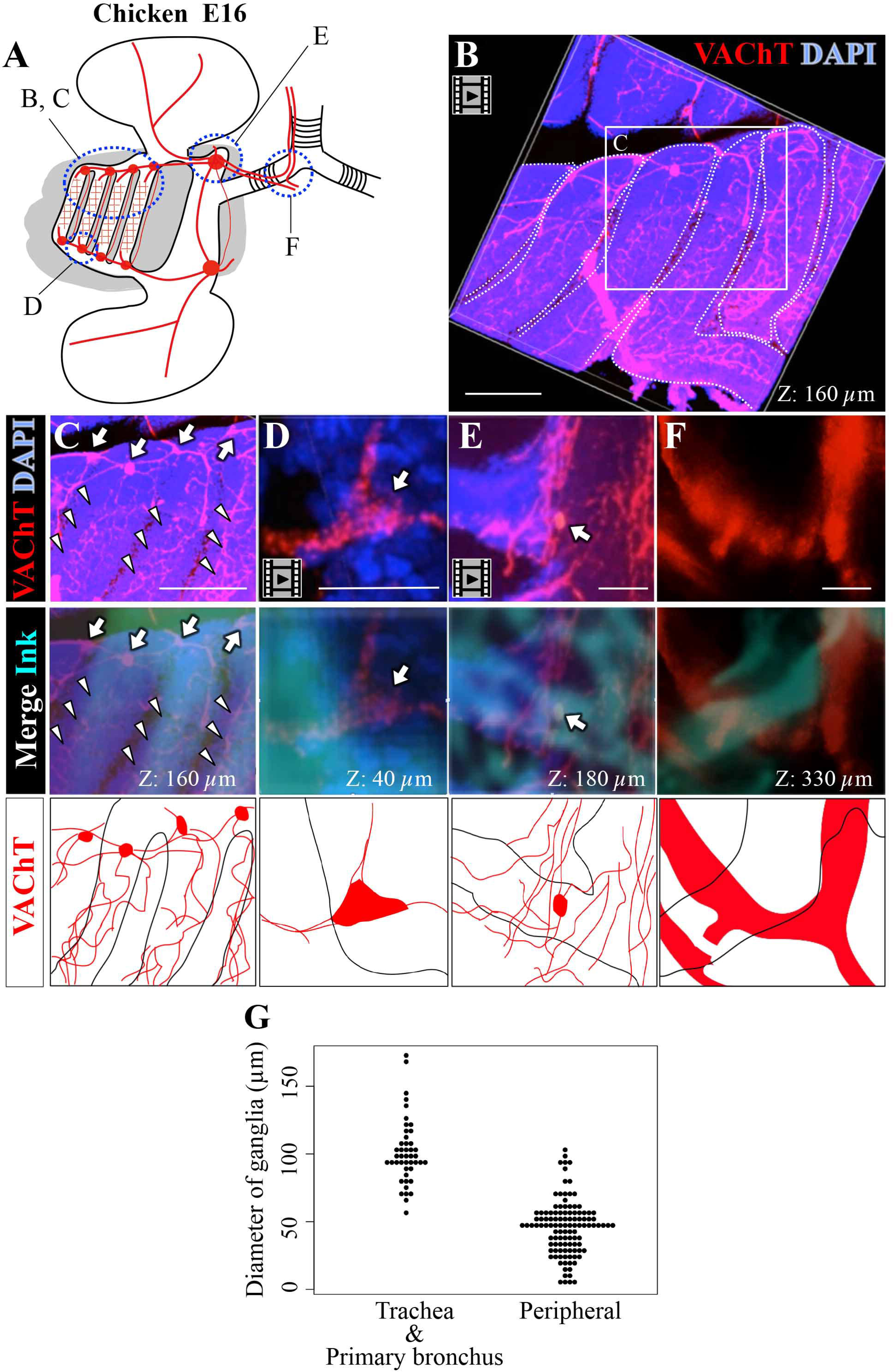
Global distribution patterns of VAChT-positive nerves and ganglia in the chicken lung at E16. (A) Diagram displaying the positions of the images (B-F), and a summary of the distribution patterns of VAChT-positive nerves and ganglia. (B-F) Confocal projection images of chicken lung at E16, stained for VAChT (red) and DAPI (blue). (B) A representative image of the movie 1 showing VAChT and DAPI signals in parabronchi. The boxed area is magnified in (C). White dotted lines encircle each bronchus. (C-F) Arrows indicate VAChT-positive ganglia. Arrowheads show mesh-like patterns of nerves in the parabronchi. Airways were visualized by ink injection (cyan). n=4 per each experiment. (G) Site-specific size of VAChT-positive ganglia in the chicken lung at E18. n=2. Confocal images were obtained at 1.0 μm intervals in (B-D), 10 μm intervals in (E-F), for Z-projections as indicated. Scale bars: 300 μm (B, C, E, F), 40 μm (D).

### Association of postsynaptic parasympathetic nerves with smooth muscle cells at the terminal bronchioles in the mouse lung

We further explored possible associations of the postsynaptic parasympathetic nerves with smooth muscles around the gas-exchanging alveoli in mice. We had initially presumed there would be a change in the patterns of VAChT-positive neurons upon the birth due to the first breath. However, the innervation patterns were essentially the same between P0^before^ and P0^after^, except for dilated bronchioles in P0^after^ (Fig. 4A-F, Movies 4, 5). Accordingly, subsequent examinations were performed either at P0^before^ or P0^after^.

**Figure 4.**
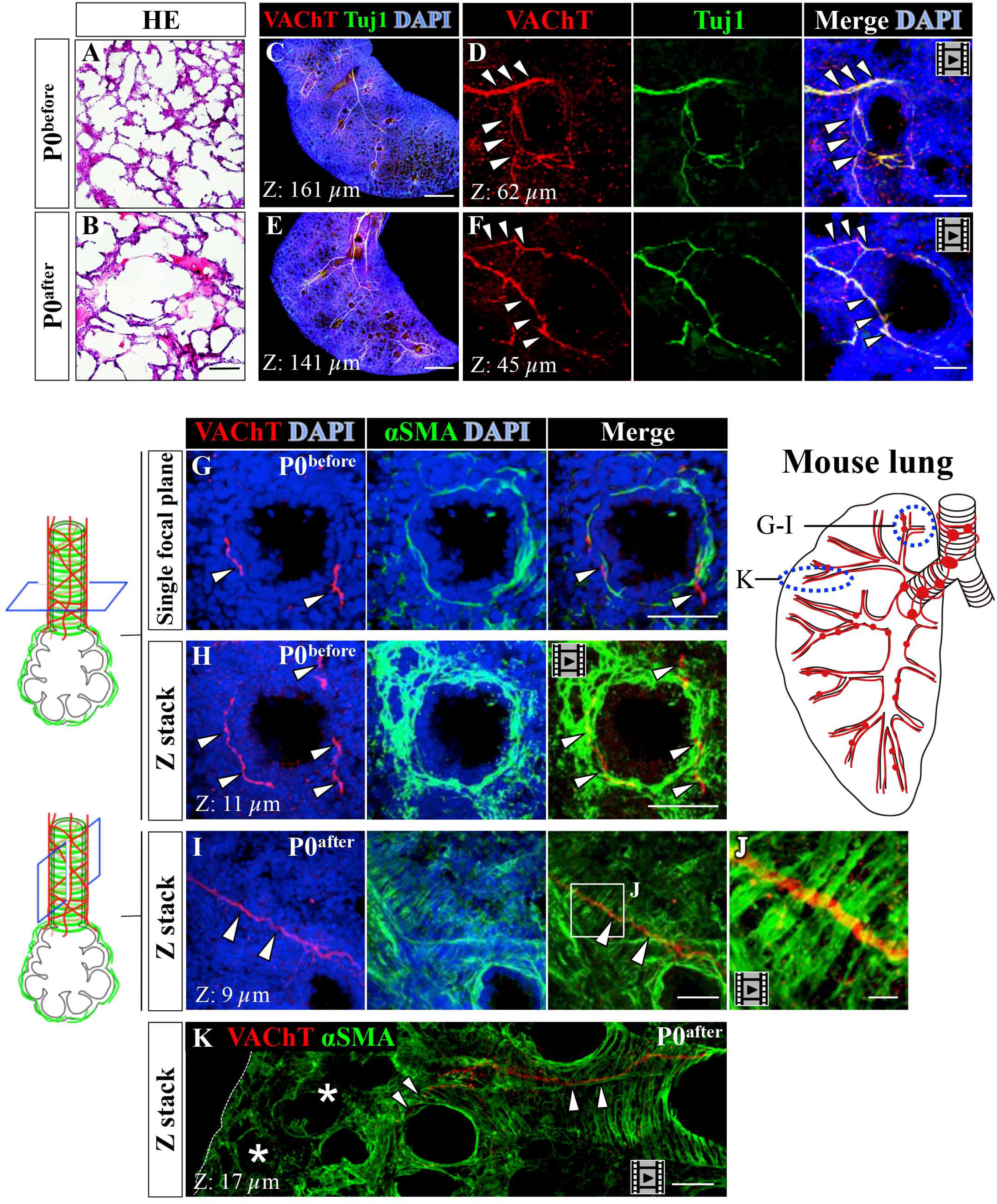
Innervation patterns of parasympathetic postganglionic nerves and smooth muscle cells in terminal bronchioles in pre- and postnatal mouse lungs. (A, B) Hematoxilyn-Eosin (HE) staining of transverse sections of the right caudal lobe at P0^before^ and P0^after^. The alveolar pore size became greater after the first breath. (C-F) Confocal projection images with staining for VAChT (red), Tuj1 (green), and DAPI (blue) of the right middle lobe. The patterns of VAChT-positive nerves remained unchanged after birth. Arrowheads indicate VAChT-positive nerves. Confocal images were obtained at 2.8 μm intervals for Z-projections as indicated. (G-K) Staining forVAChT (red), αSMA (green), and DAPI (blue) of the mouse right caudal lobe at P0^before^ and P0^after^. (G-H) Transverse sections of a terminal bronchi-containing region. Confocal images of a single focal plane (G) and Z-stack (H). (I, J) Longitudinal view of terminal bronchi. (J) is a magnified view of the square in (I). (K) Lumens of alveoli (*) and terminal bronchi can be distinguished by different patterns of αSMA-positive smooth muscles (also see text). Arrowheads indicate VAChT-positive nerves. Asterisks are alveoli. n=3 per each experiment. Confocal images were obtained at 0.15 μm intervals. Scale bars: 90 μm (A, B), 500 μm (C, E); 50 μm (D, F, G-I, K), 10 μm (J).

It is known that smooth muscle cells around terminal bronchioles are circumferentially arrayed, whereas those around alveoli are irregularly distributed (Branchfield et al., 2016). We exploited this difference as a hallmark to distinguish between alveoli and terminal bronchioles in transverse views at the periphery of the lung. By staining with anti-VAChT and anti-αSMA antibodies, we used confocal microscopy to examine transverse sections of these terminal bronchioles. As seen in a single focal plane and in the Z-stack of 11 μm, although closely associated with smooth muscle cells, VAChT-positive nerves were not circumferentially arrayed around a terminal bronchiole (Fig. 4 G, H, Movie 6). Some nerves projected in a direction parallel with the axis of the terminal bronchiole, where the nerve threaded the circumferential arrays of smooth muscle cells (Fig. 4I, J, Movie 7). Importantly, alveoli were devoid of VAChT-positive neurons (Fig. 4K, Movie 8).

### Association of postsynaptic parasympathetic nerves with smooth muscle cells in the parabronchi of the chicken lung

In chickens, gas exchange takes place in the parabronchi, where each parabronchus harbors a plethora of laterally protruding structures called atrium/atria or air capillary/capillaries. Ink injection into the airways successfully visualized a 3-D array of atria in a single parabronchus (Fig. 5A, C, E). After atria, the air goes further into air capillaries, which were not stained by the ink injection.

**Figure 5.**
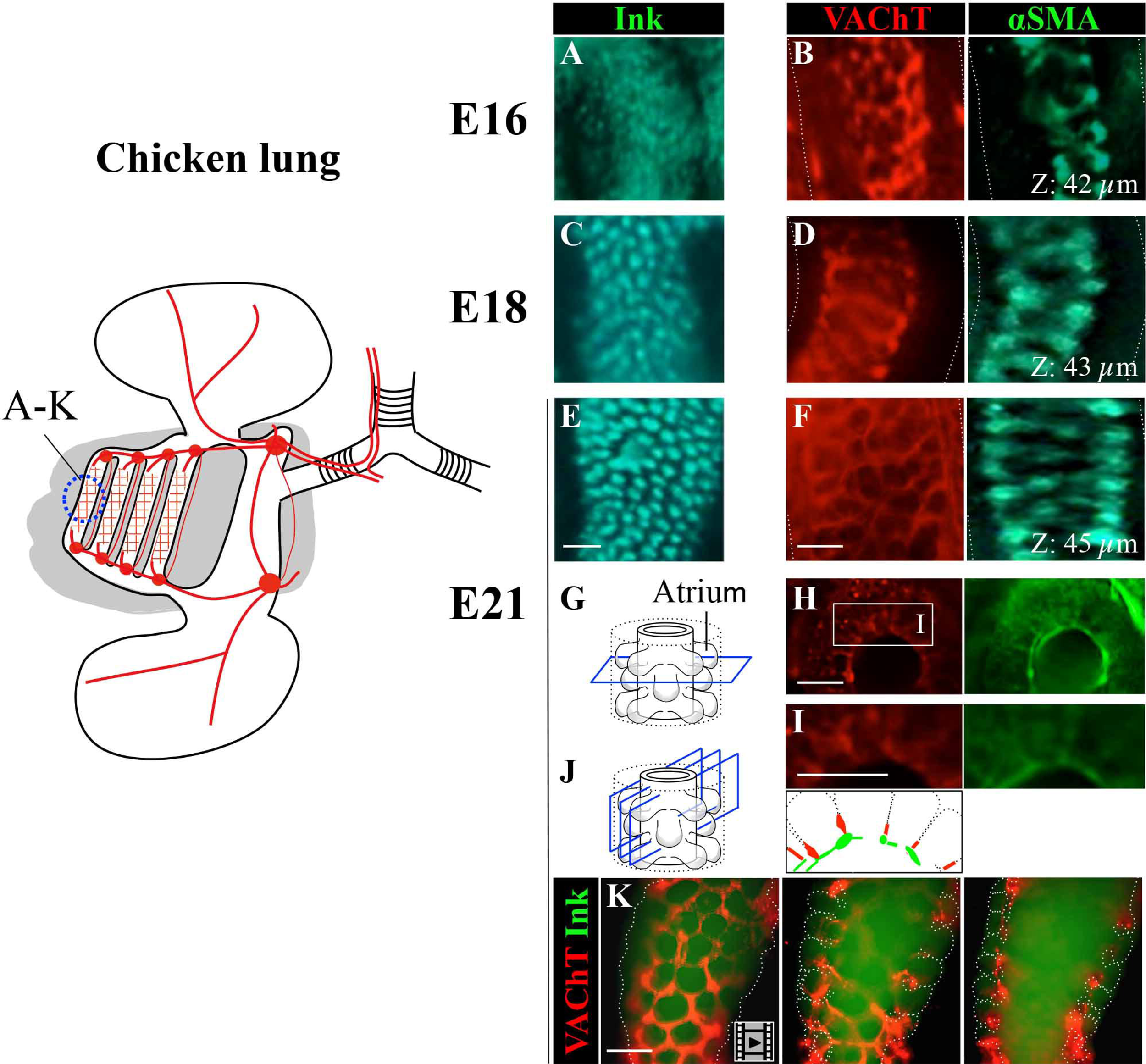
Innervation patterns of parasympathetic postganglionic nerves and smooth muscle cells in parabronchi of the chicken lung at late stages. (A-F) Projection images of parabronchial tubes from E16 to E21. (A, C, E) Longitudinal view of a parabronchus with protruding air tubes visualized by ink injection. (B, D, F) Longitudinal views of a parabronchus stained for VAChT (red) and αSMA (green). (G-I) Transverse views of a parabronchus at E21, stained for VAChT (red) and αSMA (green). (I) Magnified images of (H). (J, K) Longitudinal views of a parabronchus at E21. Nerves (VAChT) and airway (ink) are visualized. Three representative images are selected from the Movie 9. n=5 (A-F, H), n=4 (K). Confocal images were obtained at 1.0 μm intervals for Z-projections as indicated. Scale bars: 100 μm (A-K).

Parabronchi of the lung at E16 through E21 were stained with anti-VAChT- and anti-αSMA antibodies, tissue-cleared by CUBIC, and subjected to confocal microscopy. Both parasympathetic nerves and smooth muscle cells were distributed in a mesh-like pattern lining the parabronchial tubes, and such structures became progressively obvious through E21 (Fig. 5B, D, F-K, Movie 9). Notably, only the base (proximal-most portion) of each atrium was surrounded by VAChT and αSMA signals with no innervation into the gas exchange unit, which is consistent with the observation in mice (see above). Whereas smooth muscle cells were present around alveoli in mice, these cells were not observed around protruding atria or air capillaries in the chicken lung (see also Discussion).

### Sympathetic nerves in peripheral bronchi

Lastly, we compared innervation patterns of parasympathetic nerves with those of sympathetic nerves in late developing lung. It has been reported in early mouse embryos that the sympathetic nerves are located near the trachea (Lath et al, 2012), and we indeed observed Tyrosine hydroxylase (TH)-positive postsynaptic sympathetic nerves in primary and secondary bronchi in the mouse lung at P0. Double staining in sections with antibodies for TH and VAChT revealed that TH-positive or VAChT-positive neurons were either associated with each other, or present separately (Suppl. Fig. 3A-D). Of note, TH-positive neurons were also observed in peripheral regions around terminal bronchioles, although their innervation was less evident than that of parasympathetic neurons (Suppl. Fig. 3E, F, H), and some bronchioles did not harbor TH-positive nerves (data not shown). There observations are consistent with that parasympathetic nerves dominate the lung function. Like parasympathetic neurons, TH-positive nerves were not found around alveoli (Suppl. Fig. 3E, H). Intriguingly, Tuj1-positive nerves were observed in the alveoli regions, which we presume to be sensory neurons (Suppl. Fig. 3G). In chicken, Tuj1-positive nerves were sparsely located in between parabronchial tubes, showing a different pattern from that of parasympathetic nerves (Suppl. Fig. 3I, J, K).

## Discussion

We have demonstrated the 3-D innervation patterns of parasympathetic nerves and ganglia in the lungs of late developing- and newborn stages in mice and chicken embryos. The whole-mount immunostaining with anti-VAChT antibodies (Watanabe et al., 2017), the recently developed CUBIC method for tissue clearing (Susaki et al., 2014; Tainaka et al., 2014; Susaki et al., 2015), and ink-injection into airways (Takase et al., 2013) have enabled us to visualize the global structures of the parasympathetic nerves for the first time in the thick and highly elaborate lung. The parasympathetic nerves are in close association with smooth muscle cells particularly at the periphery of the lung where gas exchange units reside. Our description in the mouse lung at late stages is largely consistent with previous reports using early embryos at E12.5 that nerves innervate to the periphery of the developing lung along branching airways (Lath et al., 2012). Importantly, most of our findings obtained with the chicken lung are novel since antibodies to visualize VAChT-positive nerves in this species were not available until very recently (Watanabe et al., 2017). Furthermore, by comparing these innervation patterns between mice and chickens, remarkable similarities have emerged despite the gross difference in anatomy of the lung. The similarities include: (1) parasympathetic ganglia and nerves, presumably postganglionic fibers, are widely distributed covering the proximal and distal portions of the lung; (2) the gas exchange units, alveoli in mice and parabronchi in chickens, are devoid of parasympathetic nerves; and (3) parasympathetic nerves are in close association with smooth muscle cells, particularly at the base of the gas exchange units.

### Parasympathetic ganglia and nerves are widely distributed in the lung covering proximal and distal portions

In both mice and chickens, VAChT-positive nerves are distributed widely in the lung. Postganglionic fibers extend to the periphery along branching bronchi. Two different sizes of parasympathetic ganglia are found in a region-specific manner as summarized in Fig. 6: large ganglia reside around the first branching point near the end of trachea, whereas small ganglia are widely distributed at the periphery. Peripheral ganglia have previously been reported in mouse, pig and human embryos, although those studies did not show a region-specific localization of different size of ganglia (Sparrow et al., 1999; Weichselbaum and Sparrow, 1999; Tollet et al., 2001). The large ganglia might offer a place where vagal neurons synapse to postganglionic neural crest-derived neurons, whereas the function of small ganglia remains undetermined.

**Figure 6.**
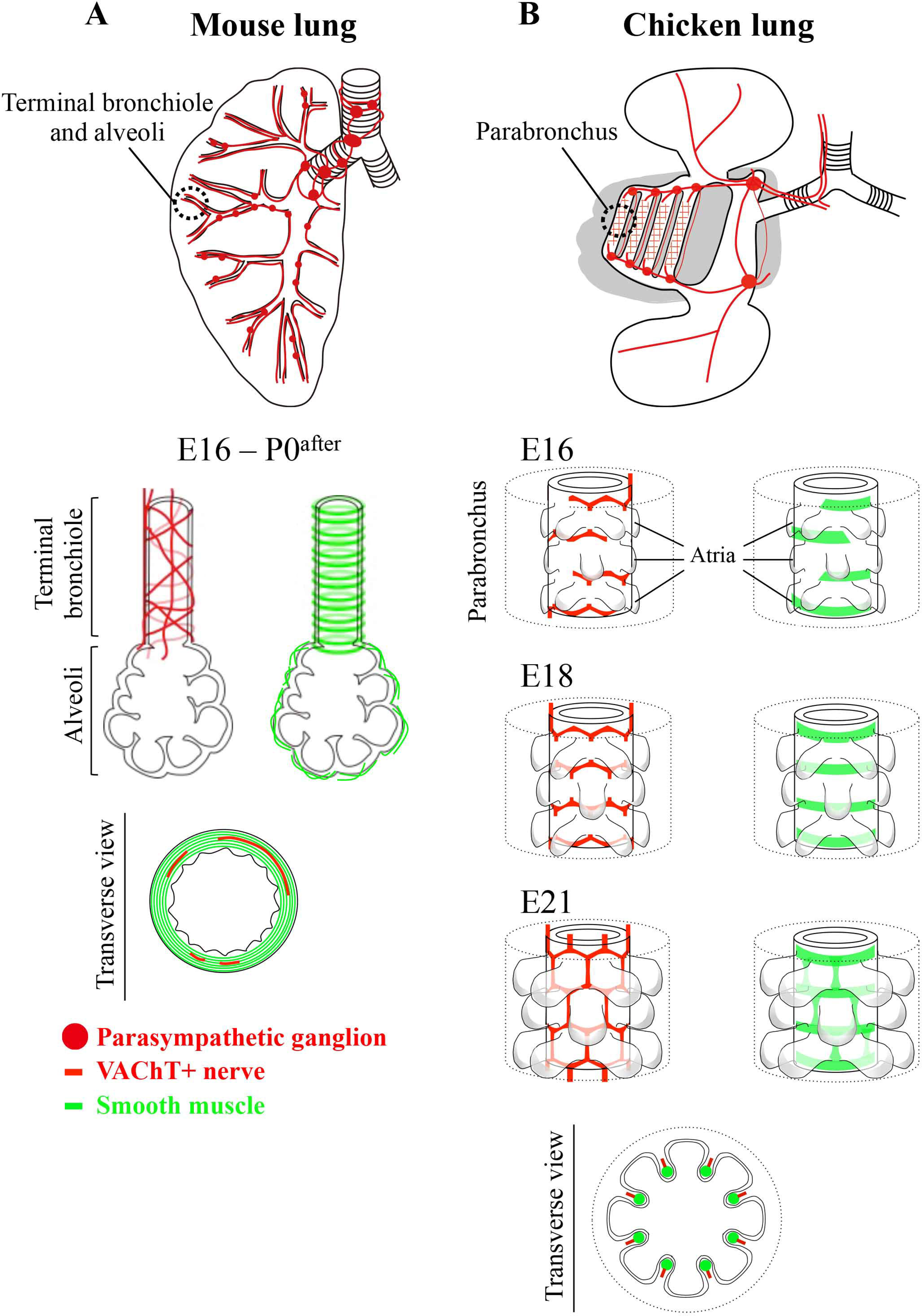
Comparison of the distribution patterns of parasympathetic nerves and smooth muscle cells in the lung between mouse (alveoli) and chicken (parabronchi). Three prominent similarities have emerged: (1) parasympathetic postganglionic fibers and ganglia are widely distributed in the lung covering the proximal and distal portions, (2) the gas exchange units, alveoli in mice and parabronchi in chickens, are devoid of parasympathetic nerves. (3) parasympathetic nerves are in close association with smooth muscle cells, particularly at the base of the gas exchange units. See also the text.

Chicken-specific distribution of ganglia should also be noted: they are found only in the main part of the lung, but not in the air sacs (Fig. 6). And the ganglia are positioned at every branching point, but not at non-branching points, e.g. within parabronchi.

### The gas exchange units, alveoli in mice and atria/air capillaries in chickens, are devoid of parasympathetic nerves

A most striking similarity in the parasympathetic innervations in the lung between mice and chickens is that the nerves do not invade into the gas exchange units, alveoli in mice and atria/air capillaries in chickens. This highlights the possibility that the gas exchange in amniotes takes place without their own constriction.

The parasympathetic innervations end at the boundary between a terminal bronchiole and alveoli in mice, and at the proximal base of each atrium in chickens. In chickens, we have succeeded in whole-mount visualization of complex arrays of atria in each bronchus, which are negative for VAChT staining (Figs. 5, 6). It is known that these atria are further connected laterally to highly intricate structures of air capillaries that also serve as a gas exchange unit (Brown et al., 1997; Makanya and Djonov, 2009; not shown in Fig. 6).

We have also demonstrated that TH-positive sympathetic nerves are absent in the gas exchange unit in both species. In mice, the TH-positive sympathetic nerves are sparsely located along terminal bronchi, showing that the dominance by the parasympathetic innervation in the lung is already established before birth.

### Parasympathetic nerves are in close association with smooth muscle cells

In both mice and chickens, close associations between parasympathetic nerves and smooth muscle cells have been found. As discussed above, parasympathetic fibers extend until the boundary between terminal bronchi/parabronchi and gas exchange units, and these innervations are accompanied by smooth muscle cells. Remarkably, in chickens, the patterns of parasympathetic fibers and smooth muscle cells are almost identical at the base of each atrium in the parabronchus (Fig. 6). During late embryogenesis, the innervation of parasympathetic fibers slightly precedes that of smooth muscle cells, suggestive of neuronal signals influencing smooth muscle cells (Fig. 6, Movie 9).

In mice, unlike chickens, the array of parasympathetic fibers in the terminal bronchioles is distinct from that of smooth muscle cells: whereas smooth muscle cells are circumferentially arrayed around the terminal bronchiole, the innervating fibers do not follow these patterns, and some nerves run perpendicularly to the smooth muscle cells (Fig. 4I, K, Fig. 6, Movies 7, 8). It has previously been reported that smooth muscle cells are also present around alveoli, but in an irregular manner, and this irregularity can be used as a hallmark to distinguish alveoli from terminal bronchioles among numerous and intricate tubular structures in the lung (Branchfield et al., 2016; Fig. 6). We have also used these criteria to locate the alveoli in αSMA–stained specimens (Fig. 4K, Movie 8). The irregularly arrayed smooth muscle cells around alveoli are not innervated by parasympathetic fibers, suggesting that these smooth muscle cells are not for constrictive regulation, but rather for physical support of the alveoli, which must be highly fragile during respiration. Collectively, it is likely that the fine-tuning of constrictive tone at the periphery of the lung is governed by coordinated functions of parasympathetic nerves and smooth muscles at the base of, but not within, the gas exchange units in both mammals and birds.

In mammals, it has previously been proposed that the parasympathetic fibers might regulate the acetylcholine-mediated constriction of smooth muscles (Sparrow et al., 1994; Sparrow et al., 1995). This notion was brought about by several separate studies. Lath et al. (2012)reported that VAChT-positive fibers were distributed along E cadherin-positive bronchioles in the embryonic lung at E12.5 and E14.5. Other studies demonstrated that nerve fibers whose identity was undetermined were in close association with smooth muscles in human and mouse fetal lungs (Sparrow et al., 1999; Tollet et al., 2001; Burns et al., 2008), and also that acetylcholine invoked bronchocontriction of a pig fetal lung *in vitro* (Sparrow et al., 1994; Sparrow et al., 1995). In the current study, we have conducted, for the first time, a direct comparison between VAChT-positive nerves and smooth muscle cells in the bronchioles/parabronchi and gas exchange units at late embryonic to postnatal stages both in mice and chickens.

### Others

In mice, the basic organization of the lung and innervation patterns of parasympathetic nerves/ganglia do not significantly change after E16, and notably, these patterns (except for the dilation of the airways) are retained after the first breath at birth (Figs. 2, 4, 6, Suppl. Fig. 1). In contrast, in chickens, the patterns of parasympathetic innervation and smooth muscle cells in the parabronchi are progressively established, and they are not yet complete at hatching (Figs. 3, 5, 6). It is conceivable that in mammals, a quick switch to the pulmonary respiration system upon the birth might necessitate the precocious establishment of the parasympathetic innervation in the lung. In contrast, a hatching chick slowly starts respiration when they are still inside the shell, and it takes about 24 hours to complete the pulmonary respiration system (Thompson, 2007).

Lastly, the comparative descriptions in this study have raised the possibility that like the mouse lung, neuroendocrine cells in the chicken lung are governed by parasympathetic nerves/ganglia. It has previously been shown in mice that clusters of neuroendocrine cells, called neuroepithelial bodies, positioned at branching points of bronchioles are innervated by VAChT-positive neurons (Domnik and Cutz, 2011; Noguchi et al., 2015). Neuroendocrine cells have also been observed in the lung of quails by electronic microscopy (Klika et al., 1998), and they are located at the “entrance into the parabronchial vestibule”, which corresponds to the transition points from a secondary bronchus to parabronchus. We have demonstrated in this study that at these transition points, VAChT-positive ganglia are preferentially located. Thus, it is conceivable that neuroendocrine cells in the lung governed by parasympathetic nervous system exert similar physiological functions between mammals and birds.

## Acknowledgements

We thank Dr. Scott F. Gilbert for helpful discussion and careful reading of the manuscript. This work was supported by JSPS KAKENHI: Grant-in-Aid for Scientific Research (B), SPIRITS (Kyoto University), and AMED (JP17gm0610015).

---

**Supplementary Figure 1**

Global distribution patterns of parasympathetic nerves and ganglia in the postnatal mouse lung at P1. (A) Confocal projection images of primary bronchus of the left lobe, stained for VAChT (red), Tuj1 (green), and DAPI (blue). (B) Confocal projection images of bronchial tubes of the left lobe, stained for VAChT (red), PGP9.5 (cyan), and DAPI (blue). Arrows indicate VAChT-positive ganglia. n=3 (A), n=2 (B). Scale bars: 100 μm. Confocal images were obtained at 3.2 μm intervals for Z-projections as indicated.

**Supplementary Figure 2**

Air sacs are poorly innervated and devoid of ganglia. (A) An anterior air sac at E21 was stained for VAChT (red) and acetylated tubulin (green). n=2. Arrowheads indicate VAChT-positive nerves. Scale bar: 300 μm.

**Supplementary Figure 3**

Distribution patterns of sympathetic nerves in postnatal lungs of mouse and chicken. (A-H) TH-staining of the mouse lung at P0 (A-G) and a diagram in the periphery (H). (A-D) Confocal projection images in the primary or secondary bronchi, stained for TH (green)/Tuj1 (red)/DAPI (blue) (A), and TH (green)/VAChT (red)/DAPI (blue) (B-D). (E-G) Confocal projection images in the peripheral region, stained for TH (green)/αSMA (red) (E), TH (green)/VAChT (red)/DAPI (blue) (F), and TH (green)/Tuj1 (red)/DAPI (blue) (G). (I-K) TH-staining of chicken lung at E21, and a diagram in a transverse view (K). A transverse section of parabronchi, stained for TH (green)/DAPI (blue) (I) and Tuj1 (red)/DAPI (blue) (J). Arrowheads in B-D and F indicate TH-positive nerves. TH-positive nerves shown by white arrowheads were closely located near VAChT-positive nerves, and TH-positive nerves shown by green ones were present singly. Asterisks are alveoli. n=3 (A-G), n=2 (I, J). Confocal images were obtained at 0.57 μm intervals in (A, G), 0.40 μm intervals in (B-D, F), 0.50 μm intervals in (E), 2.92 μm intervals in (I, J), for Z-projections as indicated. Scale bars: 50 μm (A, B, E, F, G), 25 μm (C, D), 100 μm (I, J).

**Movie 1**

Chicken lung at E16, and data source for Fig. 3B. Parabronchi were co-stained for VAChT (red) and DAPI (blue). Scale bar: 300 μm.

**Movie 2**

Chicken lung at E16, and data source for Fig. 3D. A ganglion located at the transition point from anterior bronchus to parabronchus was co-stained for VAChT (red) and DAPI (blue). Scale bar: 40 μm.

**Movie 3**

Chicken lung at E16, and data source for Fig. 3E. The transition point from the primary to secondary bronchi was co-stained for VAChT (red) and DAPI (blue). Scale bar: 300μm.

**Movie 4**

Mouse lung at P0^before^, and data source for Fig. 4C. Terminal bronchioles in the right middle lobe were co-stained for VAChT (red) Tuj1 (green), and DAPI (blue). W x H x D= 251 μm x 251 μm x 62 μm.

**Movie 5**

Mouse lung at P0^after^, and data source for Fig. 4D. Terminal bronchioles in the right middle lobe were co-stained for VAChT (red), Tuj1 (green), and DAPI (blue). W x H x D= 251 μm x 251 μm x 45 μm.

**Movie 6**

Mouse lung at P0^before^, and data source for Fig. 4H. Transverse view of a terminal bronchiole in the right caudal lobe co-stained for VAChT (red) and αSMA (green). W x H x D = 125 μm x 125 μm x 11 μm.

**Movie 7**

Mouse lung at P0^before^, and data source for Fig. 4J. Longitudinal view of terminal bronchi in the right caudal lobe co-stained for VAChT (red) and αSMA (green). W x H x D = 60 μm x 60 μm x 9 μm.

**Movie 8**

Mouse lung at P0^after^, and data source for Fig. 4K. A periphery area of the right caudal lobe containing alveoli and terminal bronchioles were stained for VAChT (red) and αSMA (green). W x H x D = 660 μm x 212 μm x 17 μm.

**Movie 9**

Chicken lung at E21, and data source for Fig. 5K. Longitudinal view of an ink-injected parabronchus stained for VAChT (red). Regularly arrayed atria protruding from the parabronchus can be seen. Scale bar: 100 μm.

## References

Balogh G., Dimitrov-Szokodi D., Husveti A., 1957. Lung Denervation in the Therapy of Intractable Bronchial Asthma. J. Thoracic Surg. 33 (2), 166–184.

Banzett, R.B., Nations, C.S., Wang, N., Fredberg, J.J., Butler, J.P., 1991. Pressure profiles show features essential to aerodynamic valving in geese. Respir. Physiol. 84 (3), 295–309.

Barnas, G.M., Mather, F.B., Fedde, M.R., 1978. Response of Avian Intrapulmonary Smooth Muscle to Changes in Carbon Dioxide Concentration. Poultry Sci. 57 (5), 1400–1407.

Boggs, D.F., Butler, P.J., Wallace, S.E., 1998. Differential air sac pressures in diving tufted ducks aythya fuligula. J. Exp. Biol. 201 (Pt 18), 2665–2668.

Branchfield, K., Li, R., Lungova, V., Verheyden, J.M., McCulley, D., Sun, X., 2016. A three-dimensional study of alveologenesis in mouse lung. Dev. Biol. 409 (2), 429–441.

Brown, R.E., Brain, J.D., Wang, N., 1997. The Avian Respiratory System: A Unique Model for Studies of Respiratory Toxicosis and for Monitoring Air Quality. Environ Health Perspect. 105 (2), 188–200.

Burns, A.J., Delalande, J.M., 2005. Neural crest cell origin for intrinsic ganglia of the developing chicken lung. Dev. Biol. 277 (1), 63–79.

Burns, A.J., Thapar, N., Barlow, A.J., 2008. Development of the neural crest-derived intrinsic innervation of the human lung. Am. J. Respir. Cell Mol. Biol. 38 (3), 269–275.

Çevik-demirkan, A., Kürtül, I., Haziroglu, R.M., 2006. Gross Morphological Features of the Lung and Air Sac in the Japanese Quail. J. Vet. Med. Sci. 68 (9), 909–913.

Cieri, R.L., Farmer, C.G., 2016. Unidirectional pulmonary airflow in vertebrates: a review of structure, function, and evolution. J. Comp. Physiol. B. 186 (5), 541–552.

Domnik, N.J., Cutz, E., 2011. Pulmonary neuroepithelial bodies as airway sensors: putative role in the generation of dyspnea. Curr. Opin. Pharmacol. 11 (3), 211–217.

Karemaker, J.M., 2017. An introduction into autonomic nervous function. Physiol. Meas. 38 (5), R89–R118.

Kitazawa, S., Fuchina, S., Suzuki, N., 1976. Studies on Air Sacs in Domestin Fowls I. Development of Air Sacs in Embryos. Obihiro University of Agriculture and Veterinary Medicine. 10, 1–24.

Klika, E., Scheuermann, D.W., De Groodt-Lasseel, M.H.A., Bazantova, I., Switka, A., 1998. An SEM and TEM study of the transition of the bronchus to the parabronchus in quail (Coturnix coturnix). Ann. Anat. 180 (4), 289–297.

Lambertz, M., Grommes, K., Kohlsdorf, T., Perry, S.F., 2015. Lungs of the first amniotes: why simple if they can be complex? Biol. Lett. 11 (1), 20140848.

Lath, N.R., Galambos, C., Rocha, A.B., Malek, M., Gittes, G.K., Potoka, D.A., 2012. Defective pulmonary innervation and autonomic imbalance in congenital diaphragmatic hernia. Am. J. Physiol. Lung Cell Mol. Physiol. 302 (4), L390–L398.

Lewis, M.J., Short, A.L., Lewis, K.E., 2006. Autonomic nervous system control of the cardiovascular and respiratory systems in asthma. Respir. Med. 100 (10), 1688–1705.

Liu, R., Song, J., Li, H., Wu, Z., Chen, H., Wu, W., Gu, L., 2014. Treatment of canine asthma by high selective vagotomy. J. Thorac. Cardiovasc. Surg. 148 (2), 683–689.

Maina, J.N., 2003. Developmental dynamics of the bronchial (airway) and air sac systems of the avian respiratory system from day 3 to day 26 of life: a scanning electron microscopic study of the domestic fowl, Gallus gallus variant domesticus. Anat. Embryol (Berl). 207 (2), 119–134.

Maina, J.N., 2008. Functional morphology of the avian respiratory system, the lung–air sac system: efficiency built on complexity. Ostrich. 79 (2), 117–132.

Maina, J.N., 2015. The design of the avian respiratory system: development, morphology and function. J. Ornithol. 156 (Suppl 1), S41–S63.

Maina, J.N., 2017. Pivotal debates and controversies on the structure and function of the avian respiratory system: setting the record straight. Biol. Rev. Camb. Philos. Soc. 92 (3), 1475–1504.

Makanya, A.N., Djonov, V., 2009. Parabronchial angioarchitecture in developing and adult chickens. J. Appl. Physiol (1985). 106 (6), 1959–1969.

Mazzone, S. B., Canning, B. J. 2013. Autonomic neural control of the airways. Handb. Clin. Neurol. 117, 215–228.

McCorry, L.K., 2007. Physiology of the Autonomic Nervous System. Am. J. Pharm. Educ. 71 (4), 78.

Metzger, R.J., Klein, O.D., Martin, G.R., Krasnow, M.A., 2008. The branching programme of mouse lung development. Nature 453 (7196), 745–750.

Nasu, T., 2005. Scanning Electron Microscopic Study on the Microarchitecture of the Vascular System in the Pigeon Lung. J. Vet. Med. Sci. 67 (10), 1071–1074.

Noguchi, M., Sumiyama, K., Morimoto, M., 2015. Directed Migration of Pulmonary Neuroendocrine Cells toward Airway Branches Organizes the Stereotypic Location of Neuroepithelial Bodies. Cell Rep. 13 (12), 2679–2686.

Plummer, E.M., Goller, F., 2008. Singing with reduced air sac volume causes uniform decrease in airflow and sound amplitude in the zebra finch. J. Exp. Biol. 211 (Pt 1), 66–78.

Reese, S., Dalamani, G., Kaspers, B., 2006. The avian lung-associated immune system: a review. Vet. Res. 37 (3), 311–324.

Sakiyama, J., Yokouchi, Y., Kuroiwa, A., 2000. Coordinated expression of Hoxb genes and signaling molecules during development of the chick respiratory tract. Dev. Biol. 227 (1), 12–27.

Schachner, E.R., Hutchinson, J.R., Farmer, C., 2013. Pulmonary anatomy in the Nile crocodile and the evolution of unidirectional airflow in Archosauria. PeerJ. 1, e60.

Sparrow, M.P., Warwick, S.P., Everett, A.W., 1995. Innervation and Function of the Distal Airways in the Developing Bronchial Tree of Fetal Pig Lung. Am. J. Respir. Cell Mol. Biol. 13 (5), 518–525.

Sparrow, M.P., Warwick, S.P., Mitchell, H.W., 1994. Foetal airway motor tone in prenatal lung development of the pig. Eur. Respir. J. 7 (8), 1416–1424.

Sparrow, M.P., Weichselbaum, M., P. B. McCray, J., 1999. Development of the Innervation and Airway Smooth Muscle in Human Fetal Lung. Am. J. Respir. Cell Mol. Biol. 20 (4), 550–560.

Susaki, E.A., Tainaka, K., Perrin, D., Kishino, F., Tawara, T., Watanabe, T.M., Yokoyama, C., Onoe, H., Eguchi, M., Yamaguchi, S., Abe, T., Kiyonari, H., Shimizu, Y., Miyawaki, A., Yokota, H., Ueda, H.R. 2014. Whole-Brain Imaging with Single-Cell Resolution Using Chemical Cocktails and Computational Analysis. Cell 157 (3),726–739.

Susaki, E.A., Tainaka, K., Perrin, D., Yukinaga, H., Kuno, A., Ueda, H.R., 2015. Advanced CUBIC protocols for whole-brain and whole-body clearing and imaging. Nat. Protoc. 10 (11), 1709–1727.

Tainaka, K., Kubota, S.I., Suyama, T.Q., Susaki, E.A., Perrin, D., Ukai-Tadenuma, M., Ukai, H., Ueda, H.R., 2014. Whole-body imaging with single-cell resolution by tissue decolorization. Cell 159 (4), 911–924.

Takase, Y., Tadokoro, R., Takahashi, Y., 2013. Low cost labeling with highlighter ink efficiently visualizes developing blood vessels in avian and mouse embryos. Dev. Growth Differ. 55 (9), 792–801.

Thompson, M.B., 2007. Comparison of the respiratory transition at birth or hatching in viviparous and oviparous amniote vertebrates. Comp. Biochem. Physiol. A Mol. Integr. Physiol. 148 (4), 755–760.

Tollet, J., Everett, A.W., Sparrow, M.P., 2001. Spatial and temporal distribution of nerves, ganglia, and smooth muscle during the early pseudoglandular stage of fetal mouse lung development. Dev. Dyn. 221 (1), 48–60.

Watanabe, T., Kiyomoto, T., Tadokoro, R., Takase, Y., Takahashi, Y., 2017. Newly raised anti-VAChT and anti-ChAT antibodies detect cholinergic cells in chicken embryos. Dev. Growth Differ. 59 (9), 677–687.

Wehrwein, E.A., Orer, H.S., Barman, S.M., 2016. Overview of the Anatomy, Physiology, and Pharmacology of the Autonomic Nervous System. Compr. Physiol. 6 (3), 1239–1278.

Weichselbaum, M., Sparrow, M.P., 1999. A Confocal Microscopic Study of the Formation of Ganglia in the Airways of Fetal Pig Lung. Am. J. Respir. Cell Mol. Biol. 21 (5), 607–620.

West, J.B., Watson, R.R., Fu, Z., 2007. The human lung: did evolution get it wrong? Eur. Respir. J. 29 (1), 11–17.

